# Spatial Transcriptomic Signature of Progressive Fibrosis in Human MASLD: Role of Senescence and Metabolic Reprogramming

**DOI:** 10.1101/2025.03.20.644239

**Authors:** Hani Vu, Yuliangzi Sun, Zherui Xiong, Daniel Radford-Smith, Andrew Causer, Christian Nefzger, Eoin O’Sullivan, Matthew Watt, Grant Ramm, Andrew Clouston, Katharine Irvine, Quan H Nguyen, Elizabeth Powell

## Abstract

Granular detail about the location and nature of liver cell interactions and the metabolic, inflammatory and fibrogenic pathways driving progressive fibrosis in metabolic dysfunction-associated steatotic liver disease (MASLD) is needed to deliver novel therapeutic targets. Here we used spatial transcriptomic data from human MASLD liver biopsies to identify the major cell types and their potential interconnected activities within specific tissue regions across the spectrum of MASLD. Gene expression data were generated using 10X Genomics Visium technology from 33 formalin-fixed paraffin-embedded liver biopsy samples and overlaid with annotated anatomical regions. Differential gene expression (DEG) and pathway analyses, cellular deconvolution and ligand-receptor interactions were conducted for each annotated anatomical category, with specific protein expression validated using CODEX spatial proteomics and immunohistochemistry staining. Unsupervised gene expression data grouped the annotated spots into 2 main clusters enriched for early/intermediate vs late fibrosis and transcriptome-based cellular deconvolution was well aligned with annotated histopathological features. In addition to extracellular matrix/receptor interactions and immune cell recruitment and trafficking, several genes encoding immunoglobulins were highly upregulated in late-stage fibrosis and were spatially associated with a senescence signature. Upregulated DEGs for early/intermediate-stage fibrosis were significantly enriched for metabolic pathways, oxidative phosphorylation and fatty acid metabolism. In contrast glycolysis genes were strongly co-expressed with late stage fibrosis. MASLD progression is accompanied by a decline in normal liver metabolic function and significant reprogramming of metabolic fuel utilisation. The spatial association of a senescence signature with expression of genes encoding immunoglobulins and complement has been linked to aging and is associated with progressive fibrosis. This work provides a valuable discovery dataset spanning different stages of human liver fibrosis and highlights the complex crosstalk between metabolic perturbations and inflammation underpinning fibrosis progression.

## Introduction

Metabolic dysfunction-associated steatotic liver disease (MASLD) is the leading cause of chronic liver disease, and a risk factor for cardiovascular events and kidney disease^1^. MASLD covers a spectrum of phenotypes, ranging from fatty (steatotic) liver without significant inflammation, to metabolic dysfunction-associated steatohepatitis (MASH) in which hepatic steatosis is associated with liver cell injury and inflammation, with or without fibrosis^2^. Patients with progressive liver fibrosis are at risk of developing cirrhosis, which in turn can lead to liver failure and hepatocellular carcinoma (HCC), a rapidly increasing form of cancer^3^.

MASLD has a complex pathophysiology: in a background of rising hepatic fat and increasing insulin resistance, inflammatory signals circulate between the liver, adipose tissue and gastrointestinal tract. These signals paradoxically increase hepatic fat production and storage, ultimately resulting in the generation of toxic lipids that drive hepatocyte damage, inflammation and fibrogenesis. As MASLD progresses, increasing fibrosis disrupts the architecture of the liver leading to cirrhosis and its inherent risk of liver failure and HCC. Key cellular mechanisms of MASLD progression include hepatocyte cell death, activation of liver-resident macrophages, immune cell infiltration, and activation of hepatic stellate cells that promote fibrogenesis^4^.

The main treatment option for people with MASLD is weight loss which is difficult to achieve and sustain with life-style modification alone^5^. Consequently, there remains an urgent need for drugs to treat the progressive form of this complex liver disease to prevent serious liver-related complications and improve cardiometabolic comorbidities. However, many compounds have failed to achieve primary endpoints in phase 3 clinical trials, likely reflecting the heterogeneity of cell types involved in the disease process as well as their variable phenotype and functional status^6^. These cell states are shaped by their location and the intercellular crosstalk within the metabolic, inflammatory and pro-fibrotic microenvironment that drives progressive MASLD.

More granularity about the location and nature of liver cell-to-cell interactions and the interconnected networks involving metabolism, inflammation and fibrosis is needed to understand the processes driving progressive fibrosis and deliver novel therapeutic targets. Advances in sequencing technology, particularly single cell RNASeq and spatial transcriptomics, provide the unprecedented opportunity to link tissue morphology with molecular profiles^7^. scRNASeq has been extensively used to generate novel insights in healthy and disease liver^8–12^, but few studies have taken advantage of spatial transcriptomics^13–15^. To our knowledge, no spatial study has been conducted to date to study fibrosis progression stages in human liver. Here we generated Visium spatial transcriptomic data from liver biopsies of patients spanning the spectrum of MASLD to identify the major cell types and their potential interconnected activities within each tissue region, and how region-specific cellular content and phenotype change with disease stage. Furthermore, we spatially characterised ligand-receptor pairs to discover region-specific cellular crosstalk associated with inflammation and fibrosis during MASLD progression.

## Results

### Spatial transcriptomic resolution of histopathological features in MASLD

Spatial transcriptomic profiling was performed on liver biopsies from 32 patients, including two from the same patient to assess the reproducibility of the experimental approach, with MASLD across the spectrum of disease severity (Figure 1A, Figure S1). Of the total 33 liver biopsies, 14 biopsies exhibited no (stage 0) or minimal (stage 1) fibrosis, 5 had intermediate (stage 2, 3a) fibrosis, and 14 displayed severe fibrosis (stage 3b, 4) (Table S1, Figure S1D). Our analysis was guided by relevant anatomical tissue regions defined by histopathological annotation and focused on changes associated with fibrosis stages (Figure 1A, 1B). The mean number of Visium capture areas (“spots”) per biopsy was 401, with a median of 1378 genes detected per spot, a common quality range for a spatial assay of archival formalin-fixed tissues, and no batch effect was observed (Figure S1A-C). Spots located on selected histopathological features were manually annotated based on the H&E-stained images of the respective sections (Figure 1B, Figure S1E). An average of > 60% of spots per biopsy were annotated. Unsupervised UMAP dimensionality reduction based on gene expression grouped the annotated spots into two main clusters (Figure 1C,D, left and right), which were enriched for spots from biopsies with late (F3b, F4) and early/intermediate (F0, F1, F2, F3a) fibrosis, respectively, demonstrating that the gene expression data contains biological information capable of differentiating fibrosis stages. Correspondingly, the late fibrosis (left) cluster contained annotated spots related to fibrous septa, portal and lobular inflammation and the portal tract interface, whereas the early/intermediate fibrosis (right) cluster contained normal and steatotic hepatocytes (Figure 1D). There was no observable difference in the distribution of ballooned hepatocytes in the late and the early/intermediate clusters (Figure 1D, 1E). There was a reduction in the proportion of spots annotated as normal hepatocytes, normal portal tracts and steatosis with advancing fibrosis stage, while no spots in the early stage biopsies were annotated as ‘fibrous septa’ (Figure 1E, Figure S1E).

**Figure 1.**
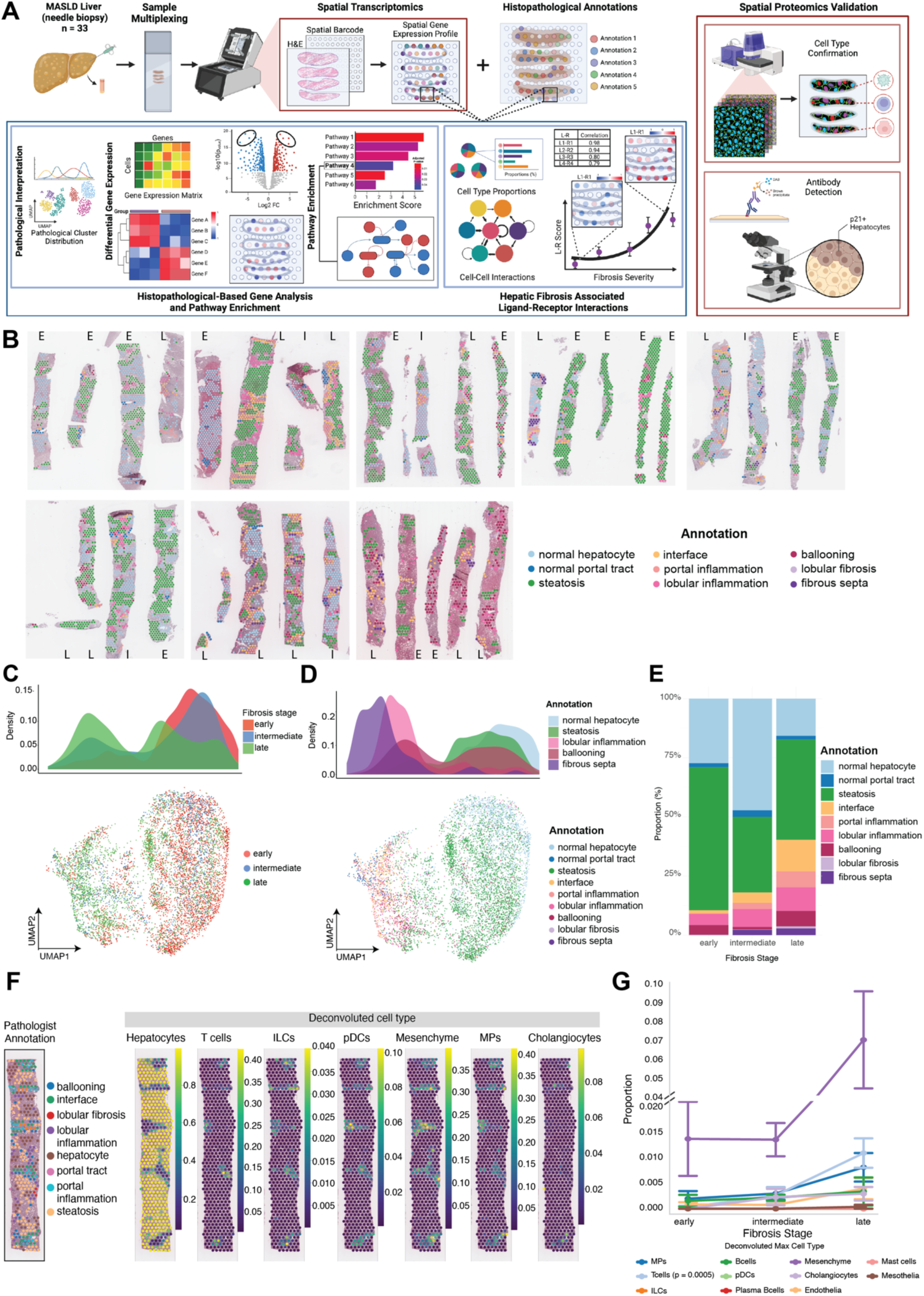
Spatial resolution of gene expression in MASLD. 33 liver biopsies from 32 patients across the spectrum of MASLD severity were profiled by Visium spatial transcriptomics. (A) Project overview highlighting the experimental workflow and focus of analysis. (B) Biopsy-overlaid Visium spots coloured by histopathological annotation and labelled by fibrosis stage; E = early, I = intermediate and L = late. Annotated spots were clustered based on gene expression using unsupervised UMAP dimensionality reduction analysis and coloured according to fibrosis stage (C) and histological annotation (D); histograms illustrate the distribution of classified spots on the X axis. (E) Mean proportion of annotated spot types among early, intermediate and late fibrosis stage biopsies. (F) Inferred deconvoluted cell type composition, displayed individually, demonstrated on a representative late-stage biopsy with scale bars representing probability. Matched biopsy with spots coloured by annotation alongside. (G) Mean proportion of deconvoluted cell types predicted by CARD analysis. Adjusted P values indicate significant linear association between cell proportion and fibrosis stage. Hepatocytes were excluded for visual clarity (see Figure S2B). MP: mononuclear phagocytes. ILC: innate lymphoid cells, DC: dendritic cells.

Cellular deconvolution based on single cell RNA sequencing from healthy and cirrhotic human liver^16^ was used to classify each spot into 12 cell types, including seven immune cell types, endothelial, epithelial (hepatocytes and cholangiocytes), mesenchymal (including hepatic stellate cells and myofibroblasts) and mesothelial cells (Figure 1F,G, Figure S2A,B). Transcriptome-based cellular deconvolution results show that the distribution of the major cell types was well aligned with annotated histopathological features (Figure 1F, Figure S2A); according to the majority cell type present. For example, steatotic regions were identified as containing a majority of hepatocyte transcripts and portal areas contained cholangiocytes, mesenchyme and inflammatory cells (Figure 1F). Although the majority of cells belong to the hepatocyte annotation category across all patients (Figure S2C), we found a significant increase in the proportion of spots attributed to mononuclear phagocytes (MPs), T cells, mesenchymal cells and cholangiocytes from early to late-stage fibrosis (Figure 1G, Figure S2B-D).

### Differential gene expression associated with specific histopathological features during MASLD progression

Genes that were significantly differentially expressed (DEGs) between early/intermediate- and late-stage fibrosis for regions annotated as steatosis, inflammation (portal and lobular combined), portal tract interface, ballooning, normal hepatocytes and normal portal tracts were identified via pseudo bulk analysis of all spatially selected spots specific for each anatomical category across all patients (Figure 2A; Figure S3A,B). Sixteen genes enriched in early- and 36 genes in late-stage fibrosis were common to steatosis, inflammation, interface and ballooning comparisons (Figure S3C,D). Portal and lobular inflammation had the highest number of DEGs compared to the other three categories.

**Figure 2.**
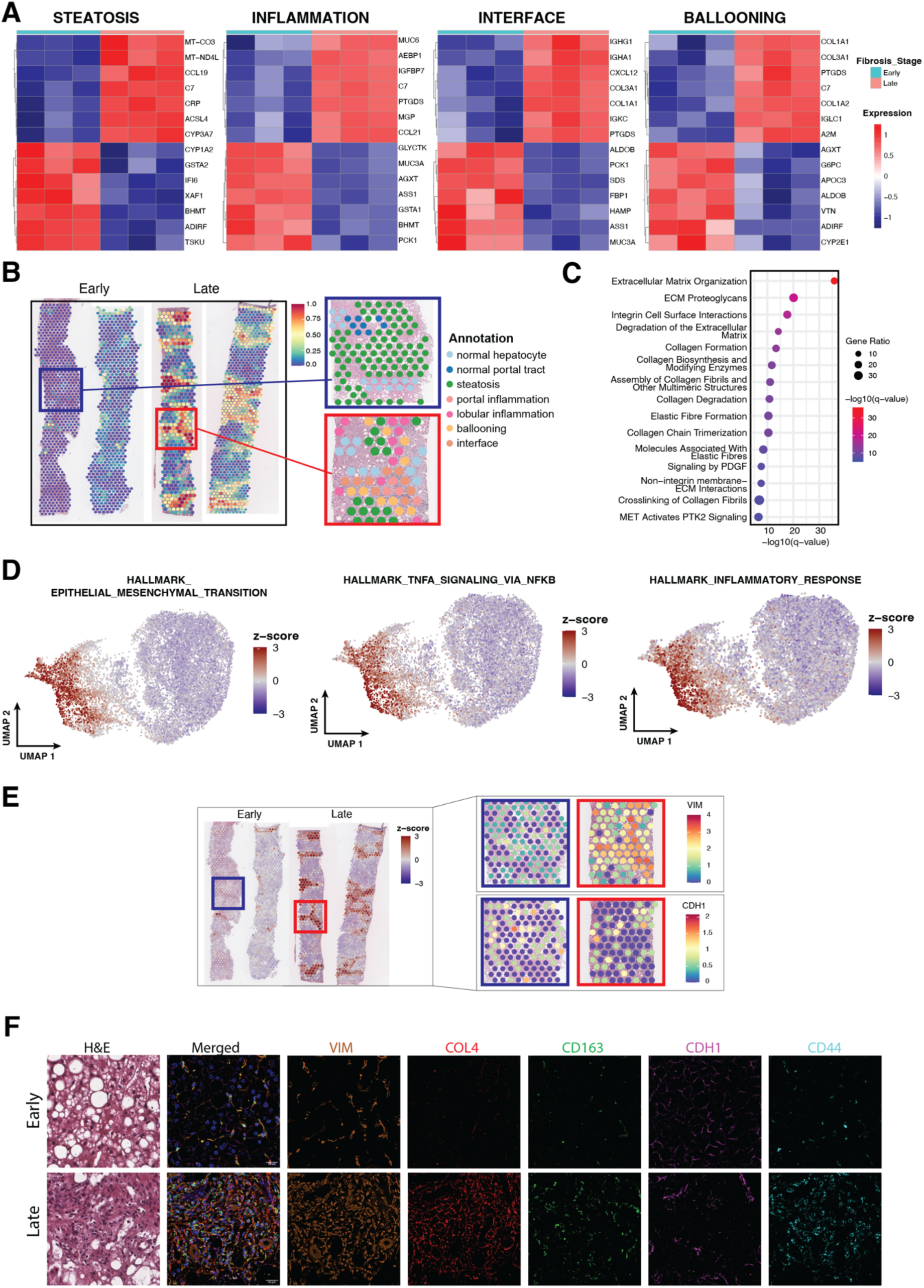
Differential gene expression associated with specific histopathological features during MASLD progression. (A) Top 7 differentially expressed genes between spots annotated as steatosis, portal or lobular inflammation, interface and ballooning in early and late-stage fibrosis. (B) Representative examples of late-stage fibrosis enriched DEGs present in two or more comparisons in early and late-stage biopsies with spots coloured by gene expression. Scale bars represent module scores of all DEG. Corresponding pathological annotations of selected regions of interest are shown. (C) Significantly enriched Reactome pathways for DEG in late-stage fibrosis. (D) Three Hallmark genesets selected from the 10 most enriched gene sets. Geneset co-expression scores were projected onto the UMAP and coloured by the scores. (E) Example spatial expression of the Epithelial-Mesenchymal Transition geneset and the top 2 related markers: vimentin (VIM) and E Cadherin (CDH1) in early and late-stage biopsies. (F) Examples of CODEX (Co-detection by indexing) stains with VIM, collagen 4 (COL4), CD163, CDH1 and CD44 protein expression detected by CODEX analysis.

Late-stage fibrosis upregulated DEGs (in two or more anatomical groups, a filter to ensure significant genes from at least two independent analyses, n = 277, Table S3) demonstrated a higher expression score in regions of portal and lobular inflammation, ballooned hepatocytes and portal interface than normal and steatotic hepatocytes (Figure 2B, Figure S4A). This gene set was enriched for multiple reactome pathways related to extracellular matrix (ECM) organisation (Figure 2C). Geneset Co-expression Analysis (GeseCA) revealed strong co-expression between UMAP-clustered late fibrosis spots and several Hallmark genesets^17^, including Epithelial-Mesenchymal Transition, TNF Signalling via NFkB, and other stress and inflammation pathways, which was clearly evident when the co-expression z-scores for each spot were projected onto the original UMAP (Figure 2D, S3E). The reduced expression of the epithelial marker E cadherin (*CDH1*) and gain of the mesenchymal marker vimentin (*VIM*) in the transition from early to late stage was spatially confirmed (Figure 2E). CODEX proteomics analysis also demonstrated higher protein expression levels of VIM and lower levels of CDH1 (Figure 2F). Collagen 4 (COL4), CD163 and CD44, which were identified as upregulated in two or more annotation groups in late fibrosis (Table S3), were also upregulated at the protein level in late-stage fibrosis (Figure 2F).

Upregulated DEGs for early/intermediate-stage fibrosis (in two or more anatomical groups, n = 304, Table S2) exhibited an inverse expression pattern compared to late-stage fibrosis DEGs in annotated regions across the liver biopsies (Figure 2B, Figure 3A). These genes were significantly enriched for metabolic pathways, including amino acids, lipids, fatty acids, bile salts and xenobiotics (Figure 3B). GeseCA also revealed strong co-expression of UMAP-clustered early stage fibrosis spots and these physiological metabolic pathways (Figure S3E). Glycolysis genes were strongly co-expressed with late stage, whereas genes involved in oxidative phosphorylation and fatty acid metabolism were enriched in early stage fibrosis spots (Figure 3C). The latter, along with the significant enrichment in the Biological Oxidations pathway among early stage upregulated genes (Figure 3B), may reflect impaired mitochondrial function and fatty acid oxidation associated with MASLD progression^18^. This apparent metabolic reprogramming may also indicate progressive erosion of hepatic metabolic zonation, as has previously been suggested^19,20^. Oxygen-demanding processes such as gluconeogenesis, beta-oxidation and ammonia conversion to urea are higher in zone 1 (periportal), whereas glycolysis, lipid synthesis and ammonia conversion to glutamate are higher in zone 3 (pericentral) hepatocytes^19^. Consistent with portal erosion, expression of the liver isoform of glutaminase (GLS2) and rate-limiting enzyme of the urea cycle (CPS1) were increased in early-stage biopsies, whereas glutamine synthetase (GLUL) was upregulated in late-stage fibrosis (Table S2-3). Gluconeogenic genes FBP1, encoding the rate-limiting enzyme Fructose 1,6 bis-Phosphatase 1, PCK1, ALDOB, G6PC and ASS1, responsible for metabolism of the urea cycle and glucogenic amino acid arginine^21^, were similarly enriched in early-stage fibrosis (Table S2). The switch from reliance on oxidative phosphorylation to glycolysis, known as the Warburg effect, is also frequently reported in other inflammatory states and cancer. Consistent with this, metabolic genes found to be upregulated in HCC^22^ were enriched in late-stage fibrosis (Figure 3D,E), whereas HCC-downregulated metabolic genes^22^ were enriched in the early-stage cluster, demonstrating lower expression in portal and lobular inflammatory regions (Figure 3F,G). A meta-analysis of transcriptomic data identified over 600 metabolic genes that were consistently up- or down-regulated in human hepatocellular carcinoma samples^22^. Visualisation of spatial gene expression changes for all genes as well as highly enriched metabolic pathways are shown in Figure S4 and Figure S5. We further analysed a lipidomic dataset for an extended cohort of 218 patient samples with fibrosis from early to late stages.

**Figure 3.**
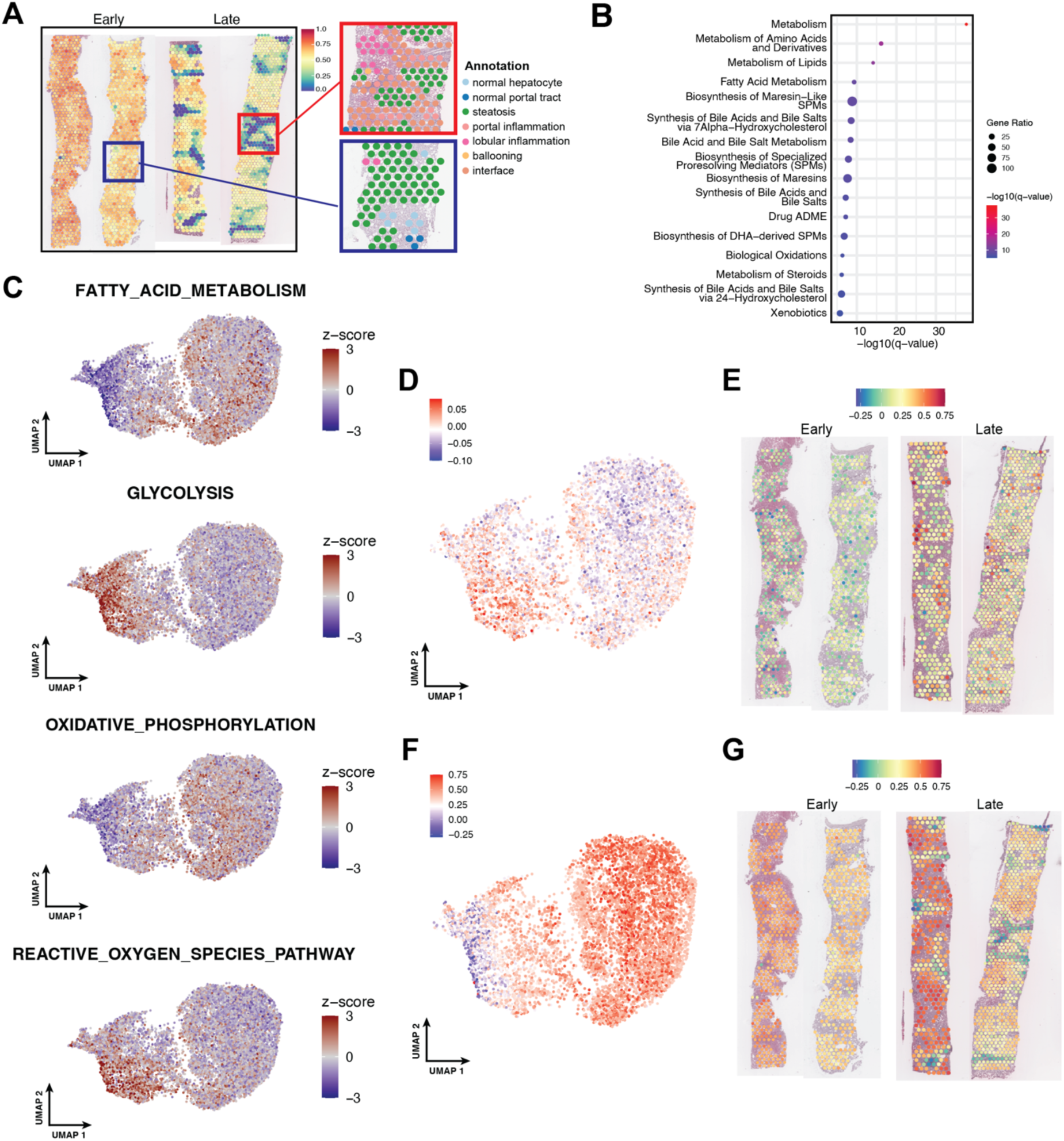
MASLD Progression is associated with metabolic reprogramming at a transcriptional level. (A) Representative examples of early-stage fibrosis enriched DEGs present in two or more annotomical comparisons between early and late-stage biopsies with spots coloured by gene expression. Scale bars represent module scores of all DEGs. Corresponding pathological annotations of selected regions of interest are shown. (B) Significantly enriched Reactome pathways for DEG in early-stage fibrosis. (C) Geneset co-expression scores for the metabolic-related hallmark gene set projected onto the UMAP. Expression of metabolic genes identified as up- (D, E) or down-regulated (F, G) in HCC for each spot visualised by UMAP clustering (D,F) and biopsy location (E,G).

Among the top five annotated metabolites that changed, we found a reduction of Fumaric acid, a metabolite important in the tricarboxylic acid (TCA) energy metabolism (Figure S6). Taken together, these molecular signatures are consistent with the increase in matrix deposition and inflammatory signalling with MASLD progression, a concomitant decline in normal liver metabolic function, and significant reprogramming of metabolic fuel utilisation.

### MASLD progression is associated with increased immunoglobulin expression and senescence

Several genes encoding immunoglobulins (*IGKC, IGHG, IGHA, IGLC*) were highly upregulated in late-stage fibrosis at the portal tract interface and in regions containing ballooned hepatocytes (Figure 2A). Ma et al (2024) recently reported immunoglobulin and complement expression co-localised with accumulating senescence ‘hotspots’ in aged tissues, including mouse and human liver^23^. Cellular senescence is also a feature of chronic liver disease, including MASLD^24,25^. We therefore hypothesised that immunoglobulin gene expression would be spatially associated with a senescence signature in MASLD biopsies. Differentially expressed genes between spots with a high immunoglobulin module score (comprising *IGKC, IGHG, IGHA, IGLC*), compared to those with a low module score, included multiple genes previously implicated in Senescence-Associated Secretory Programs (SASPs) in various tissues, including *TGFB1*, *TIMP2*, *MMP2*, *IGFBP7* and *CXCL12*^26–28^ (Figure 4A). *CDKN1A*, encoding the canonical senescence-associated cell cycle inhibitor p21, was upregulated in late fibrosis (Table S3), and significantly increased at the protein level in hepatocyte nuclei in late stage biopsies (Figure 4B, C). We created an integrated senescence signature based on the SenMayo^26^ and Gene Ontology Bioprocess genesets (Table S5). The senescence signature was abundantly enriched in inflammatory and fibrotic regions when projected onto the original UMAP and represented spatially (Figure 4D,E). Consistent with an association between senescence and immunoglobulin expression, there was a positive correlation between the senescence gene module score and the immunoglobulin module score within each spot, with the highest correlation in the fibrous septa and lobular fibrosis (Figure 4F). Consistently, the mean senescence module score was highest in fibrous septa, along with portal inflammation (Figure 4G). Despite the increase in immunoglobulin expression, our cell deconvolution analysis did not detect any enrichment of plasma cells, likely due to the relatively small proportion of these cells (Figure 1G), consistent with the previous report^23^. Senescence in MASLD has also previously been linked with similar hepatic metabolic perturbations to those identified by our Differential Gene Expression and GeseCA analyses above and the development of HCC, with the gluconeogenic gene FBP1 increasingly implicated as an important tumor suppressor. Taken together, these data suggest increased immunoglobulin expression and cellular senescence is associated with MASLD progression.

**Figure 4.**
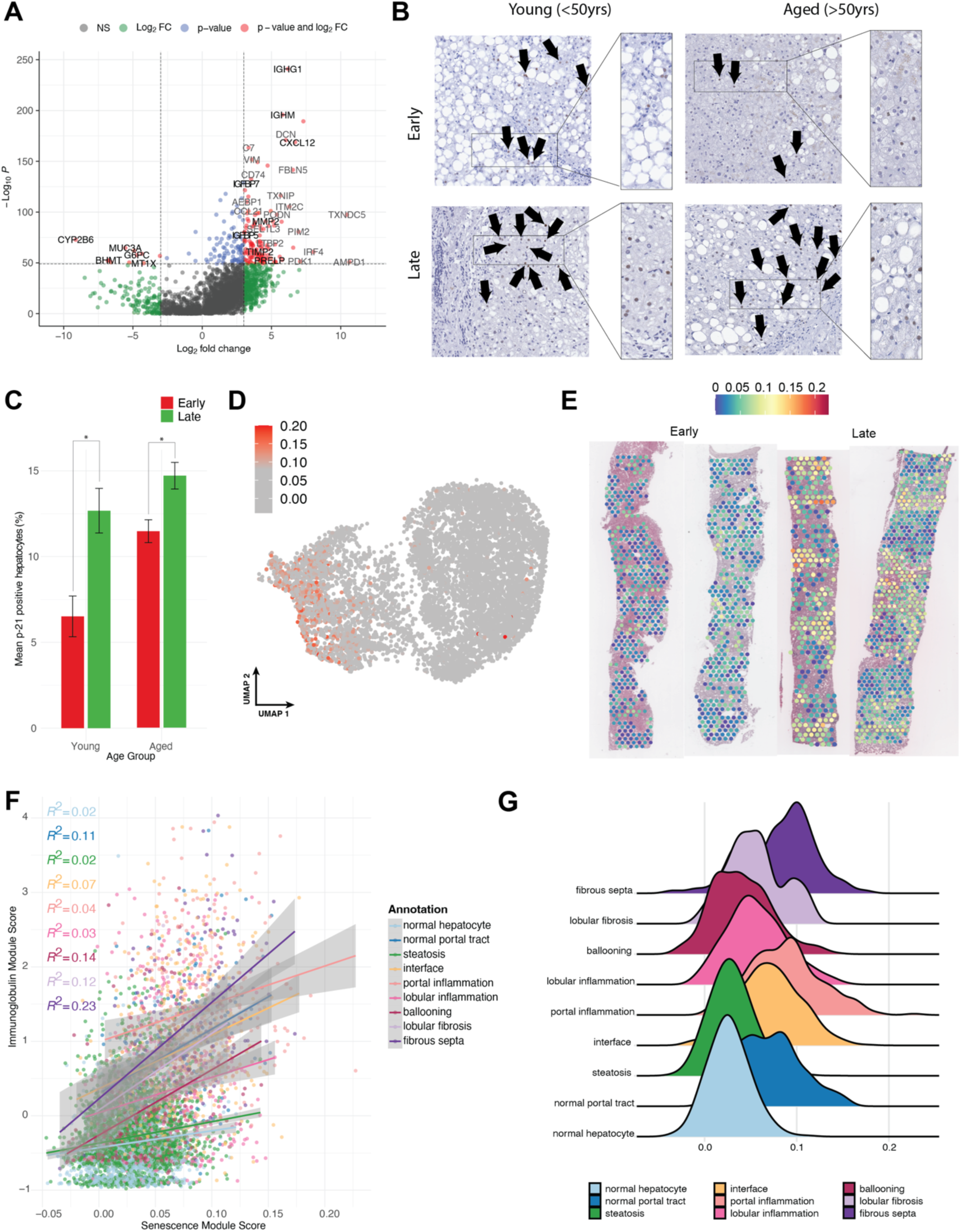
MASLD progression is associated with increasing cellular senescence. **(A)** Differentially expressed genes between spots with the highest and lowest predicted immunoglobulin expression. (B) Immunohistochemistry staining for p21 on representitve young (<50 years) and aged (>50 years) samples for early and late fibrosis stage. (C) Bar plot showing mean percentage of p21 positive hepatocytes per biopsy with error bars showing standard deviation. Student’s t-test, *P≤0.05. Senescence gene expression score plotted spatially on representative early- and late-stage biopsies. (D) Spearman correlation between the average expression of the genes in the senescence module and immunoglobulin module score for each spot (E). Ridge plot showing mean module score for expression of senescence genes, grouped by annotation

### Co-regulated Ligand-Receptor pairs associated with MASLD fibrosis progression

To gain further insight into the cellular and molecular interactions driving fibrosis progression in MASLD, we applied an unbiased, high-throughput screening approach to identify ligand-receptor pairs (LRPs) that were statistically enriched for spatial co-expression within Visium spots, using spatially-constrained two-level permutation (SCTP) test implemented in stLearn^29^. To ensure comprehensiveness, we utilised connectomeDB2020 (consisting of 2293 manually curated ligand-receptor pairs with literature support)^30^. Ligand-receptor scores between neighbouring cells, reflecting the probability and strength of interactions, were tested for significant correlation with fibrosis stages, suggesting increasing or decreasing interactions with progression from early to late fibrosis (Figure 5A,B). Fibrosis associated LRPs included extracellular matrix components (collagens interacting with CD44, CD36 and DDR2, decorin (DCN) and biglycan (BGN) interacting with TLR2) and modulators (eg. Tissue Inhibitor of Metalloproteinase 1 (TIMP1)-CD63) and inflammatory chemokines, cytokines and receptors (eg SPP1-ITGAV, CXCL12-CD4, CLEC11A-ITGB1) (Figure 5C,D). Of those, the predicted interaction between TIMP1-CD63 was most strongly correlated with fibrosis progression (Figure 5C,D; Figure S7A), consistent with the previous report that this interaction activates the PI3K/Akt survival pathways, preventing apoptosis of activated fibroblasts and hepatic stellate cells, which drive fibrosis^31–33^. Examples of LRPs with higher activity in early disease included RTN4 and its receptor, ANGPTL3-ITGAV, EpherinA1 and its receptors EPHA1 and EPHA2 (Figure 5C,E; Figure S7). The LRP analysis was then confined to portal inflammation annotated spots, to identify candidate LRPs driving inflammation in this key region. The LRPs most highly correlated with fibrosis stage included inflammatory mediators such as CXCL12-ACKR3, SEM4AD-PLXNB2 and LTB-LTBR (Figure 5F). Several LRP in portal inflammation spots were inversely correlated with fibrosis, including signalling pathways such as C3-CR1, MIF-CD74, and LGALS9-CD44 (Figure 5G). The chemokine CXCL12 was a component of nine of the 145 LR pairs associated with increasing fibrosis stage, including seven of the top 50. Although CD4 was the most highly correlated candidate receptor, CXCL12’s canonical signalling receptors CXCR4 and CXCR7 (ACKR3) were also identified, along with several integrins (Figure 5C). CXCL12 is one of a number of chemokines that have been reported as part of the senescence associated secretory program (SASP)^27^, though not specifically in the liver. We identified DEGs between spots with highest and lowest scores for CXCL12 interaction with any putative receptor (top and bottom 10%, respectively). Genes associated with high CXCL12-Receptor activity included other SASP genes such as IGFBP7^34^ and CCL5, along with immunoglobulins and the known pro-fibrotic cytokine osteopontin (SPP1) (Figure 5H).

**Figure 5.**
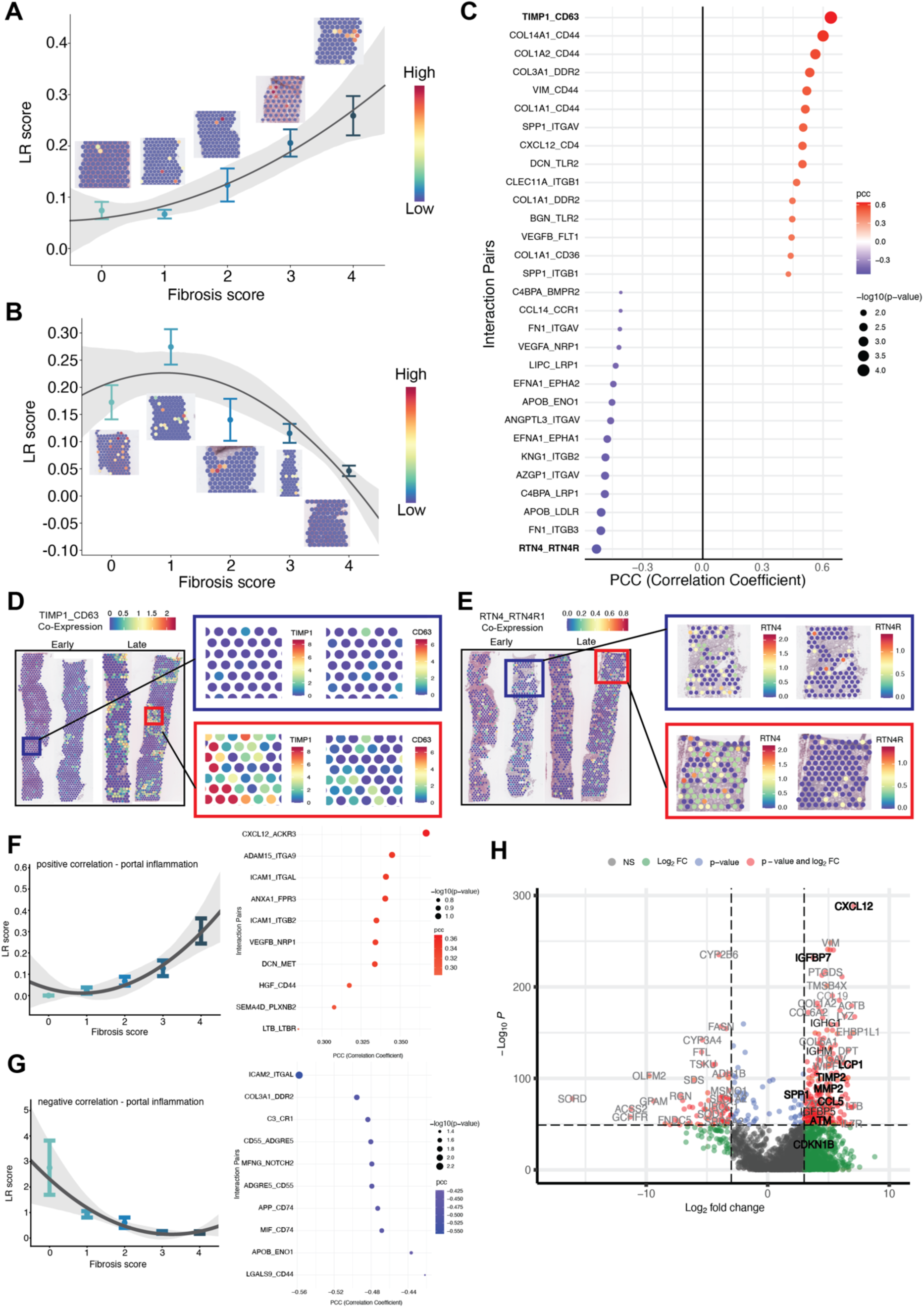
Co-regulated Ligand-Receptor pairs associated with MASLD fibrosis progression. Ligand-receptor (LR) pair interactions were predicted based on co-expression by neighbouring cells within spots and assigned an LR score based on the strength of the prediction. LR pairs positively (A) or negatively (B) correlated with fibrosis stage were identified. Top 15 LR pairs, correlation shown by gradient scale from blue to red and -log(p-value) represented by the size of dots (C). Spatial visualisation of the top LR pairs positively (TIMP1-CD63) and negatively (RTN4-RTN4R1) correlated with fibrosis stage (D,E). LR pair interactions were predicted based on co-expression within spots annotated as portal inflammation and assigned an LR score based on the strength of the prediction. LR pairs positively (F) or negatively (G) correlated with fibrosis stage were identified. Differentially expressed genes between spots with the highest and lowest predicted CXCL12-Receptor activity for all identified interacting receptors (H).

## Discussion

The development of progressive fibrosis in MASLD involves dynamic cellular interactions within the highly structured microenvironment of hepatic lobules. The liver’s complex organisation and pleiotropic metabolic functions have led to difficulty in achieving effective drug therapies for MASLD fibrosis. In this study we demonstrated that MASLD progression is associated with upregulation of a senescence gene expression signature and p21 expression in hepatocytes, associated with increased immunoglobulin gene expression, and concomitant perturbation of hepatic metabolic pathways.

Not surprisingly, as fibrosis progressed there was an overall significant increase in mesenchyme, mononuclear phagocytes, T cells and cholangiocytes, with an upregulation in genes enriched for extracellular matrix/receptor interactions (MGP, COL3A1, COL1A1, COL1A2), and immune cell recruitment and trafficking (CCL19, CCL21, CXCL12). In particular, these transcripts were among the most highly differentially expressed genes in the portal tract interface and regions containing ballooned hepatocytes that release inflammatory and fibrogenic signals^35^. Although many of these DEGs have previously been reported in progressive MASLD, our spatial transcriptomics data provide novel information about their location and the associated cellular heterogeneity. Gene sets highly correlated with genes regulated during MASLD progression include TNF Signalling, reflecting the strong inflammatory basis of the process and EMT, previously proposed as a possible driver of liver fibrosis^36,37^. We^24,38^ and others^39^ identified that the ductular reaction (DR), a complex of strings of cholangiocytes and bile ductules, in a niche of stromal and inflammatory cells at the portal tract interface, correlates with the extent of fibrosis in MASLD. As well as producing factors that lead to increased extracellular matrix, ductular cells share some properties of mesenchymal cells^40^, although whether they are generated via EMT remains unclear^41,42^.

As fibrosis progressed, the differential expression of ligand-receptor signalling pairs revealed cell-cell communication networks dominated by extracellular matrix components/modulators and inflammatory chemokines, cytokines and receptors. These interactions included recognised drivers of fibrosis such as osteopontin^43,44^, vimentin^45^ and CXCL12^46^, and illustrate the redundancy in inflammatory and fibrogenic signalling^47^. The chemokine CXCL12 featured in nine interactions within ligand-receptor pairs that correlated with fibrosis severity (CD4, ITGA4, ITGAV, ITGB1, CXCR4, ITGA5, ACKR3, ITGA5, ITGB3), with different interactions dominating in specific histopathological regions. The CXCL12 signalling axis has a role in the migration and activation of leucocytes, endothelial and stem cells, via binding to CXCR4 (leading to G protein signaling) and the scavenging receptor ACKR3 (that signals via the β-arrestin-2 pathway). Mouse models of chronic liver injury suggest that the ACKR3 and CXCR4 pathways have differential effects on liver repair and fibrosis respectively. Inhibiting the CXCL12 – CXCR4 pathway did not reduce fibrosis^46^, whereas blockade of CXCR4-mediated inflammation counteracted age-related senescence and steatosis and mitigated diet-induced MASLD^48^. More recently, CXCL12 has been reported to activate integrins in a rapid, receptor-independent manner^49^, suggesting a role in modulating leucocyte extravasation. Our data also support the relevance of this chemokine in human MASLD, and the need for more detailed investigations into the role/s of CXCL12/CXCR4 signaling in liver fibrosis.

Our study provides further insight into the possible *causal* factors that may be driving this marked inflammatory and fibrogenic response. Unexpectedly, genes encoding immunoglobulins (*IGKC, IGHG1, IGHA1, IGLC1*) were among the top eight genes differentially expressed between early and late-stage fibrosis at the portal tract interface and in regions containing ballooned hepatocytes. Although not previously reported in MASLD, immunoglobulin-related genes were the most highly upregulated DEGs in aged tissues from male mice and located near senescent cells^23^. IgG accumulated in adipose tissue in aging mice, activating a macrophage inflammatory response leading to adipose tissue fibrosis and impairing metabolic health^50–52^. In our human MASLD liver biopsies, the UMAP cluster related to fibrous septa, inflammation and the portal interface showed enrichment in genes and pathways including *TGFB1*, *TIMP2*, *MMP2*, *IGFBP7* and *CXCL12* that are linked to senescence pathways. The accumulation of senescent cells in MASLD contributes to liver disease progression via activation of the senescence-associated secretory phenotype (SASP)^27,53^. Many of the upregulated genes in our inflammatory/fibrotic cluster are recognised components of the SASP, and include chemokines such as CXCL12 discussed above, growth factors and proteases that impact on the surrounding tissue microenvironment^27,54,55^. In mice, elimination of senescent cells reduced hepatic steatosis^56^ but senolytic drugs did not reduce MASLD progression^57^. It is likely that targeting senotherapeutic strategies towards distinct senescent cell types will be important to remove or counteract their effect in specific contexts, in order to reduce liver injury and improve immunosurveillance^58^.

Our data show marked dysregulation in metabolic pathways with increasing fibrosis, particularly in regions defined by inflammation, ballooning and at the portal tract interface. Several of the genes expressed at low levels in late stages of fibrosis (ALDOB, FBP1, PCK1, BHMT, GSTA1, ASS1, AGXT, GLYCTK, G6PC) likely reflect the metabolic adaptations in cellular substrate metabolism required for activation of inflammatory and fibrogenic cells^22,59–63^. Increased aerobic glycolysis is a key feature of activated hepatic stellate cells and inflammatory macrophages and preclinical studies have shown these cells are also influenced by hepatocyte metabolic changes^60,63^. For example, hepatocyte-specific loss of the gluconeogenic enzyme fructose 1,6-bisphosphatase 1 (FBP1, one of our top ‘down-regulated’ genes between early- and late-stage fibrosis at the portal tract interface) promoted stellate cell activation and senescence in rodent models^60^. FBP1 is increasingly implicated as a tumor suppressor gene. FBP1 was reported to be universally silenced in human and murine liver tumours, and loss of hepatocyte FBP1 in mice promoted tumorigenesis via promoting hepatic stellate cell senescence^60^. Another recent report identified FBP1 as a p53 target, which was elevated in pre-malignant senescent hepatocytes, but re-activated in malignant hepatocytes to enhance activity of previously senescent cells, thereby facilitating carcinogenesis^64^. Disease-linked metabolic remodeling may in part be set up by predictable age-related tissue changes, as evidenced by the rising incidence of MASLD from the fourth decade of life onward. Reported age-associated tissue alterations include shifts in liver zonation in both mice and humans^23^, including downregulation of CDH1, a periportal zonation marker. We also identified CDH1 downregulation along with reduced expression of other periportal genes (GLS2, CPS1, gluconeogenic genes including FBP1), with concomitant upregulation of the pericentral GLUL. Age-related metabolic programming may thus be one mechanism contributing to the increased risk of MASLD with advancing age, raising the possibility that MASLD may, in part represent an accelerated form of liver aging. Targeting these metabolic adaptations that drive inflammatory and fibrogenic cellular responses may have therapeutic benefit in MASLD^63^, however cell-type and context-specific targeting of metabolic pathways will likely be crucial to minimise off-target events.

Our unbiased spatial analysis provides an integrated view of molecular pathway alterations and cellular interactions spanning the spectrum of MASLD. We identify metabolic perturbations and increasing senescence and inflammatory pathways that are significantly linked to fibrosis progression. These candidate molecules warrant validation through independent cohorts and functional perturbation models.

## Methodology

### Ethics approval and Tissue Collection

This study was approved by the QIMR Berghofer Human Research Ethics Committee (Project No. 3924) and Metro South Health Governance (Approval Number SSA/2023/QMS/104139). Formalin-fixed paraffin-embedded (FFPE) liver biopsy samples were available for 32 patients with MASLD who were recruited to a study investigating risk stratification of MASLD^65^. Sex and age of patients, as well as severity of fibrosis were considered during sample selection to provide a spectrum of disease severity and clinical characteristics. Liver histology was assessed by an experienced liver pathologist (A.D.C.) who was blinded to the clinical and transcriptomic data. Fibrosis stage was recorded using a modified Non-Alcoholic Steatohepatitis Clinical Research Network (NASH CRN) staging score^66^ based on the extent of steatosis, inflammation, hepatocellular ballooning and fibrosis. Fibrosis was staged from 0 to 4 as follows: stage 1, zone 3 perisinusoidal only or portal/periportal only; stage 2, zone 3 perisinusoidal and portal fibrosis; stage 3, bridging fibrosis; and stage 4, cirrhosis. Stage 3 - bridging fibrosis – was subdivided into stage 3a (few [1 or 2] fibrous septa) and stage 3b (many [>2] fibrous septa), which has been shown to give a more uniform spread of fibrosis stage^67^. Fibrosis was then categorised as early (stages 0 and 1), intermediate (stages 2 and 3a) or late (stages 3b and 4). (Figure 1B, Figure S1A,B).

### Sample Preparation

FFPE liver needle biopsies tissue sections (5um thickness) for each patient were provided on regular (uncharged) microscopic slides. Histopathological assessment was conducted on an adjacent slide to determine a tissue region of interest for spatial measurement. Four to five needle biopsies were detached from the original uncharged slides and transferred into a 6.5mm x 6.5mm region on a single SuperFrost charged slide. This way the samples were multiplexed in a nested design so that each spatial slide had samples from more than one fibrosis stages to avoid technical batch effect. Using Methyl-acrylate, we detached the tissues from their original slides, lifted and transferred the tissue to the SuperFrost charged slide for spatial measurement. This microarray slide was then utilised for spatial transcriptomics library preparation.

### Spatial Transcriptomics

#### Library Preparation

Each slide was processed following the 10X Genomics Demonstrated Protocol (#CG000520) in preparation for downstream library construction. Deparaffinization was performed, followed by Hematoxylin staining (Mayer’s Haematoxylin Solution, Sigma) for 5 minutes and counterstaining with Eosin (Alcoholic Eosin Y Solution, Sigma) for 1 minute. High-resolution brightfield imaging was conducted using the Leica Aperio XT Automated Slide Scanner at 40X magnification. Following imaging, each array was de-stained using 0.1N HCl and de-crosslinking was performed with TE buffer (pH 7) at 70°C for 1 hour.

#### Library Construction and Sequencing

Spatial transcriptomic libraries were prepared using the Visium CytAssist Spatial Gene Expression for FFPE, Human Transcriptome, 6.5mm kit (10X Genomics, PN #1000520) according to the Visium Spatial Gene Expression User Guide Rev C (10X Genomics, #CG000495). Amplification of the final Visium libraries was performed using 13–17 PCR cycles, followed by qualitative assessment.

Paired-end dual-indexed sequencing was performed on the NovaSeq 6000 platform (Illumina) using the following read configuration: Read1 – 28 bp, i7 – 10 bp, i5 – 10 bp, Read2 – 50 bp. Raw sequencing base call (BCL) data were demultiplexed using the Bcl2Fastq conversion software (v2.20).

### Image Alignment and Gene Expression Mapping

Alignment of gene expression data with corresponding high-resolution Hematoxylin and Eosin images was performed manually using Loupe Browser (10X Genomics, v7.0). Slide layout files were obtained from 10X Genomics to ensure precise spatial registration. Sequencing reads were aligned to the GRCh38 human reference genome using the SpaceRanger computational pipeline (10X Genomics, v2.0). Spatial barcode information was utilized to map gene expression data to the corresponding coordinates on the histological images.

### Manual Pathological Annotations

The .cloupe file output from SpaceRanger (10X Genomics, V2.0), was loaded into the LoupeBrowser 6 user interface to explore processed data. Anatomical regions across each liver biopsy were assessed and annotated by A.D.C. Regions annotated included portal tracts with and without portal inflammation, the portal tract interface (to include areas of periportal ductular reaction), areas of lobular inflammation, normal hepatocytes, steatotic hepatocytes, ballooned hepatocytes, lobular fibrosis, and fibrous septa.

### Quality Control and Normalization of Visium Data

Mapped and aligned data were processed using the Seurat R package (v5.1.0) for downstream analysis. Gene expression levels were quantified based on unique molecular identifier (UMI) counts. Outliers and low-quality reads were removed by filtering spots with fewer than 10 counts per spot and sparse genes expressed in fewer than three spots. Additionally, spots with mitochondrial gene content exceeding 10% were eliminated to ensure data quality.

The filtered data were normalised and scaled using Seurat’s SCTransform function, which employs a regularised negative binomial regression to retain biological variability while reducing technical artifacts across sequencing runs^68^. The 3,000 most variable genes identified through this process were selected for downstream dimensionality reduction and clustering analyses.

### Integration and Dimensionality Reduction

To correct for batch effects, data integration was performed using Seurat’s IntegrateData function. The top 3,000 variably expressed genes from each sample were used to compute anchors via Canonical Correlation Analysis (CCA) and Mutual Nearest Neighbors (MNN) analysis, thereby reducing technical variability^69^.

Dimensionality reduction was achieved through Principal Component Analysis (PCA), with the first 30 principal components (PCs) selected based on elbow plot visualization. Uniform Manifold Approximation and Projection (UMAP) embeddings were then generated to facilitate visualization, with spots labeled according to histological annotations and fibrosis stages.

### Pseudo-Bulking and Differential Gene Expression Analysis

To compare gene expression between early/intermediate- and late-stage fibrosis, pseudo-bulk analysis was performed by aggregating gene expression counts across three randomly generated pseudo-replicate groups per fibrosis stage. Using the aggregateAcrossCells function from the scater package, raw UMI counts for each gene were summed within each cluster^70^. Genes with low expression in the pseudo-bulked clusters were excluded, and read counts were normalized using the filterByExpr and calcNormFactors functions from the edgeR package^71^.

Differential gene expression (DEG) analysis was conducted separately for each annotated anatomical category using the edgeR framework. Covariates were accounted for using the limma package, providing robust statistical inference^72^. A negative binomial model was fitted using glmQLFit to correct for overdispersion, and differentially expressed genes were identified using glmTreat based on a false discovery rate (FDR)-adjusted p-value < 0.05 and a log fold change (logFC) threshold of 1.2. Heatmaps generated with the pheatmap package were used to visualize gene expression patterns across each group. Genes consistently upregulated each fibrosis group in at least two anatomical categories were retained for downstream gene set enrichment analysis.

### Gene Ontology (GO) Analysis of DEGs

To identify biological processes associated with the DEGs in each annotation category, Gene Ontology (GO) enrichment analysis was performed using the ClusterProfiler package. The enrichGO function, along with the human genome database, was used to determine significantly enriched GO terms (p < 0.01). The identified terms were grouped into broader ontology tags and visualised using REVIGO^73^.

### Gene-Based Co-Regulation Pathway Analysis

Pathway co-regulation analysis was conducted using the Hallmark gene set collection from the Molecular Signatures Database (MSigDB). Gene Set Co-expression Analysis (GeseCA), implemented in the fgsea package, was applied to assess co-variation of genes within predefined pathways in early- and late-stage fibrosis. Utilising PCA loadings, pathway-level co-expression scores were computed and were then statistically tested using permutation-based null distributions, which were generated by randomly selecting gene sets of the same size. The co-regulation scores were mapped onto UMAP embeddings using the plotCoregulationProfileReduction function in fgsea. The top three hallmark gene sets for both early- and late-stage fibrosis were identified and used for further analysis.

### Module scores

Module scores for a given list of genes were calculated using the AddModuleScore function implemented in Seurat. The score was calculated based on the average expression of genes in the list subtracted to the background score, which was computed as the average of control genes randomly selected from the same dataset.

### Deconvolution

The composition of cell types within each Visium spot was estimated using Conditional autoregressive-based deconvolution (CARD) using the human fibrotic liver single-cell reference reported by Ramachandran et al (2019)^16^. The 12 lineage annotations from the original publication were utilised. Changes in cell type proportions from early and late-stage fibrosis samples were examined.

### Ligand-Receptor (L-R) Interactions

We applied stLearn spatially-constrained two-level permutation (SCTP) method to infer and analyse intercellular communication networks in Visium data^29^. SCTP first identifies spatial neighborhoods of L-R co-expression, computing so-called LR scores for each individual Visium spot, followed by a unique constrained, two-level permutation test of both genes and spots to robustly identify spatial locations where a given L-R pair has significantly higher scores than random. The L-R score for each Visium spot can be plotted onto the tissue.

#### Finding L-R pairs associated with fibrosis stages

Fibrosis scores, with ordinal values ranging from 0 to 4, were used to calculate Pearson correlation coefficient (pcc) with L-R scores for each of the significant L-R pairs across all spots in the dataset. The p-value for the correlation was calculated for significantly different to 0. L-R pairs were ranked by correlation values, p-values, and mean expression of the L-R across all spots where they coexpressed.

#### Downstream analysis of L-R interaction

To test for possible effects of L-R interaction to gene expression at the spatial locations where the interactions were predicted to occur, we categorised spatial spots into two groups of high vs. low expression of the L-R. In this study, we focused on CXCL12, which was found to be most common among the L-R highly correlated with fibrosis stages. The CXCL12 high spots were then compared with low spots for identifying differentially expressed genes, which were visualised by a volcano plot.

### Immunohistochemistry (IHC) Staining

Four 5um thick liver biopsy samples (2 x early-stage fibrosis and 2 x late-stage fibrosis) were deparaffinised using the Leica Autostainer XL . Heat induced antigen retrieval was carried out in a pressure cooker at 100°C for 20 minutes, followed by a 1% H_2_O_2_ treatment at room temperature for 10 minutes. Sections were incubated overnight at 4°C with the primary mouse monoclonal antibody for p21 (Merck Millipore, #OP64), at 1:50 dilution in the Background Sniper (Biocare Medical, SKU: BS966) blocking buffer. Sections were incubated with the MACH 2 Universal HRP-Polymer (Biocare Medical, SJU: M2U522) secondary antibody for 30 minutes at room temperature. Finally, a hematoxylin counterstain was performed using the Leica Autostainer XL, including serial dehydration with alcohol, then xylene before coverslipping.

p21-positive hepatocytes were quantified using QuPath (v 0.5.1) with the positive cell detection measurement parameter. Nuclear segmentation was applied with an intensity threshold of 0.2, while a further threshold of 0.1 was used to identify p21-positive nuclei. To reduce false positives and exclude non-hepatocyte structures, nuclei with a circularity below 0.65 were filtered out. Liver biopsies were split into three areas and the mean percentage of p21-positive hepatocytes was calculated, with error bars showing standard deviation. As p21 is also a marker of cellular aging, patient age at the time of biopsy was considered. A t-test was performed to assess statistical differences between early and late-stage biopsies within young (<50 years) and aged (>50 years) age groups.

### Spatial Proteomics - PhenoCycler

Multiplexed imaging was performed using the CODEX (CO-Detection by Indexing) system (Akoya Biosciences) on formalin-fixed paraffin-embedded (FFPE) liver needle biopsy tissues. Three 5um FFPE liver biopsy samples (1 x early stage fibrosis, 1 x intermediate stage and 1x late stage fibrosis) were mounted onto poly-L-lysine-coated coverslips, deparaffinized, rehydrated, and subjected to heat antigen retrieval using Dako Epitope Retrieval Buffer (pH 9.0) in a Biocare Medical Decloaking Chamber.

For antibody staining, sections on coverslips were incubated overnight at 4°C with a 35-plex antibody panel (Table S4), pre-conjugated with unique DNA barcodes. Following post-staining fixation, iterative reporter hybridization and imaging cycles were performed using a Zeiss Axioscope microscope. The imaging process involved sequential hybridization of fluorescently labeled oligonucleotide reporters, followed by imaging and signal removal before the next cycle. Acquired images were processed and analyzed using the CIM (CODEX Instrument Manager) and Akoya Biosciences analysis pipeline.

### Lipidomics analysis

The patient cohort is described in Patel et al. (2018)^65^. A total of 218 samples were grouped into fibrosis stages based on the clinical diagnosis of the advanced fibrosis system, with scores ranging from 0 (no evidence of fibrosis from biopsy) to 5 (evidence of fibrosis from both biopsy and imaging). Three fibrosis groups were defined: Early (scores 0, 1), Intermediate (scores 2, 3), and Late (scores 4, 5). The raw data were log-transformed after adding 10^-6^ and subsequently normalized by total signal, ensuring that the total signal for each patient matched the geometric mean of all samples. After normalization, Z-transformation was applied to each metabolite across all patients. Differential abundance analysis was performed on the normalized, transformed data using a two-sided Welch’s t-test, which does not assume equal variance. The effect size was quantified using Cohen’s d to measure the difference between group means.

