## Supplementary Figures for "Spatial Transcriptomic Signature of Progressive Fibrosis in Human MASLD: Role of Senescence and Metabolic Reprogramming"

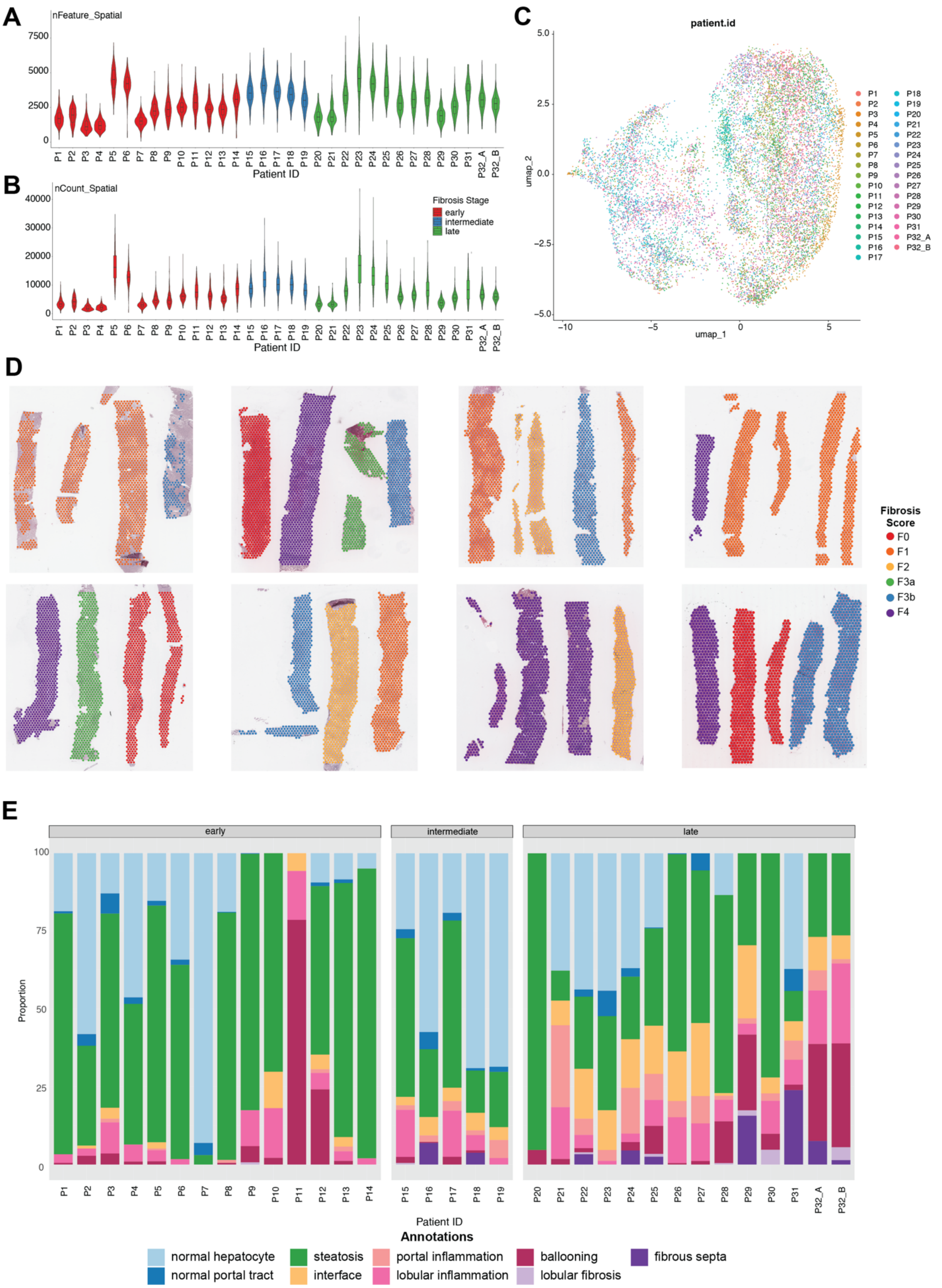

**Figure S1. Overview of the spatial transcriptomics data and pathological annotations across liver biopsies.** Scatter plot showing (A) number of genes (nFeature) and (B) total number of genes (nCount) detected per Visium spot for each liver biopsy, grouped by fibrosis stage. (C) UMAP plot demonstrating integration of all annotated spots across liver biopsies of all patients, demonstrating no observable batch effect. (D) Spatial visualisation of all liver biopsies overlayed with annotated Visium spots, coloured according to fibrosis score. (E) Proportion of annotated regions for each patient, grouped by fibrosis stage.

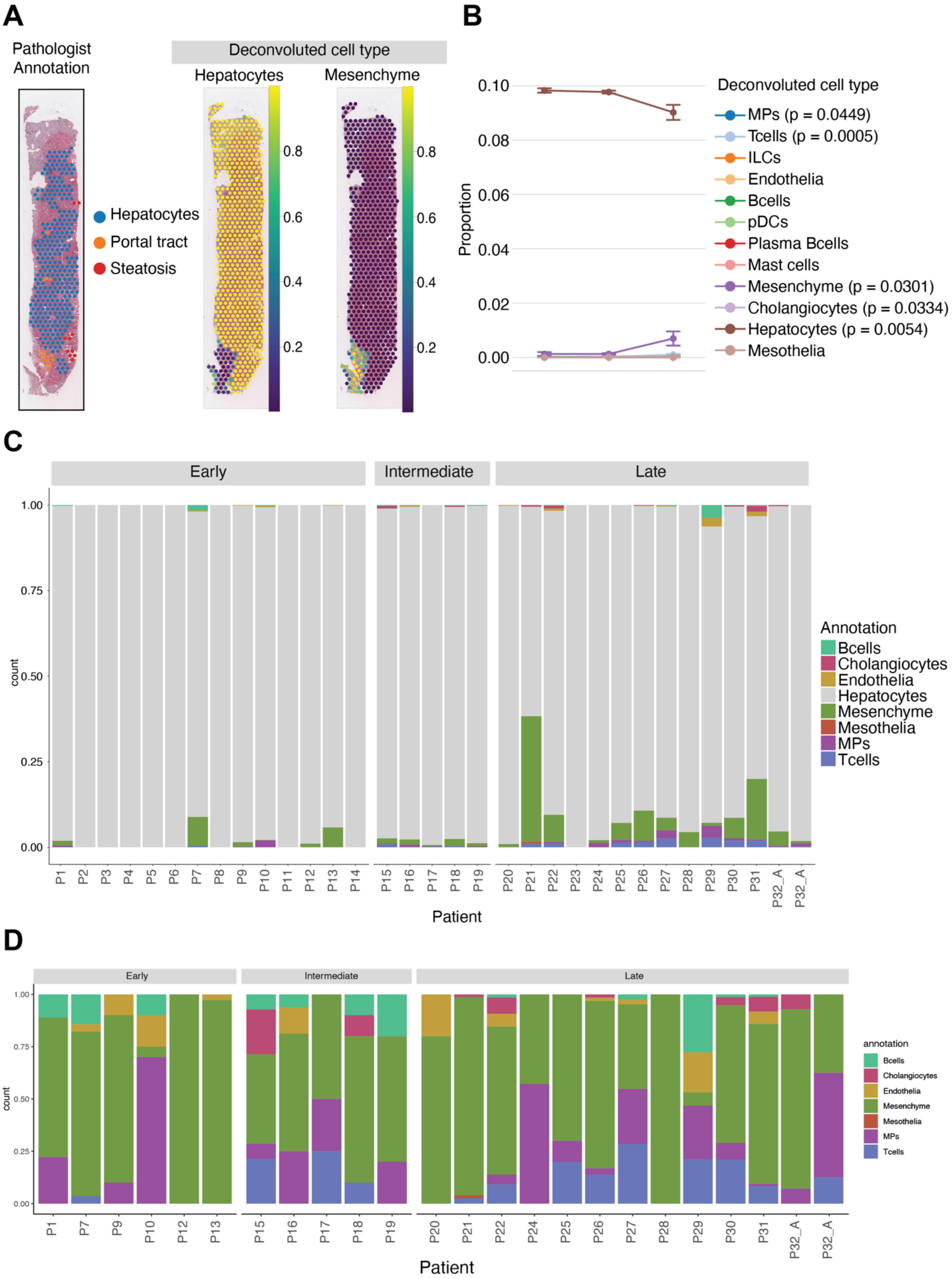
A
Pathologist Annotation
Deconvoluted cell type
Hepatocytes
Mesenchyme
Hepatocytes
Portal tract
Steatosis
0.8
0.6
0.4
0.2
B
Proportion
0.10
0.08
0.06
0.04
0.02
0.00
Deconvoluted cell type
MPs (p = 0.0449)
Tcells (p = 0.0005)
ILCs
Endothelia
Bcells
pDCs
Plasma Bcells
Mast cells
Mesenchyme (p = 0.0301)
Cholangiocytes (p = 0.0334)
Hepatocytes (p = 0.0054)
Mesothelia
C
Early
Intermediate
Late
count
1.00
0.75
0.50
0.25
0.00
P1 P2 P3 P4 P5 P6 P7 P8 P9 P10 P11 P12 P13 P14 P15 P16 P17 P18 P19 P20 P21 P22 P23 P24 P25 P26 P27 P28 P29 P30 P31 P32_A P32_A
Patient
Annotation
Bcells
Cholangiocytes
Endothelia
Hepatocytes
Mesenchyme
Mesothelia
MPs
Tcells
D
Early
Intermediate
Late
count
P1 P7 P9 P10 P12 P13 P15 P16 P17 P18 P19 P20 P21 P22 P24 P25 P26 P27 P28 P29 P30 P31 P32_A P32_A
Patient
annotation
Bcells
Cholangiocytes
Endothelia
Mesenchyme
Mesothelia
MPs
Tcells

**Figure S2 Cell type deconvolution across fibrosis stage.** (A) Representative early-stage fibrosis liver biopsy, comparing pathological annotation with deconvolution results. (B) Deconvoluted cell type proportion across fibrosis stage, including hepatocyte abundance. (C) Distribution of deconvoluted cell types across patients, including hepatocyte proportions. (D) Distribution of deconvoluted cell types across patients, excluding hepatocytes.

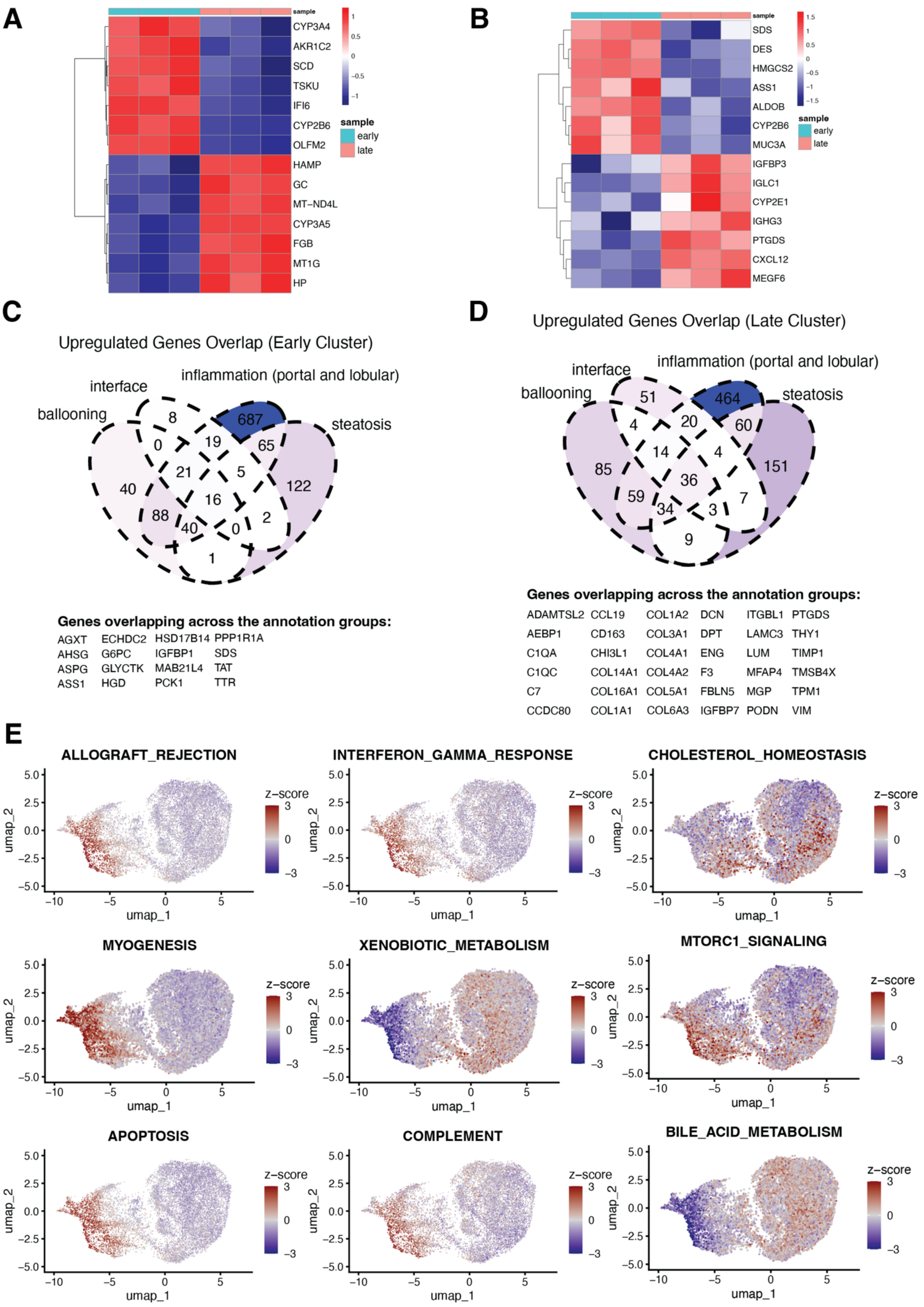

A
sample
CYP3A4
AKR1C2
SCD
TSKU
IFI6
CYP2B6
OLFM2
HAMP
GC
MT−ND4L
CYP3A5
FGB
MT1G
HP
early
late
B
sample
SDS
DES
HMGCS2
ASS1
ALDOB
CYP2B6
MUC3A
IGFBP3
IGLC1
CYP2E1
IGHG3
PTGDS
CXCL12
MEGF6
early
late
C
Upregulated Genes Overlap (Early Cluster)
interface
inflammation (portal and lobular)
ballooning
steatosis
8
687
0
19
65
21
5
40
122
16
88
2
40
0
1
Genes overlapping across the annotation groups:
AGXT ECHDC2 HSD17B14 PPP1R1A
AHSG G6PC IGFBP1 SDS
ASPG GLYCTK MAB21L4 TAT
ASS1 HGD PCK1 TTR
D
Upregulated Genes Overlap (Late Cluster)
interface
inflammation (portal and lobular)
ballooning
steatosis
51
464
4
20
60
14
4
85
36
151
59
34
3
7
9
Genes overlapping across the annotation groups:
ADAMTSL2 CCL19 COL1A2 DCN ITGBL1 PTGDS
AEBP1 CD163 COL3A1 DPT LAMC3 THY1
C1QA CHI3L1 COL4A1 ENG LUM TIMP1
C1QC COL14A1 COL4A2 F3 MFAP4 TMSB4X
C7 COL16A1 COL5A1 FBLN5 MGP TPM1
CCDC80 COL1A1 COL6A3 IGFBP7 PODN VIM
E
ALLOGRAFT_REJECTION
INTERFERON_GAMMA_RESPONSE
CHOLESTEROL_HOMEOSTASIS
MYOGENESIS
XENOBIOTIC_METABOLISM
MTORC1_SIGNALING
APOPTOSIS
COMPLEMENT
BILE_ACID_METABOLISM
umap_1
umap_2
z−score

**Figure S3. Differential gene expression (DEGs) and gene set co-expression analysis across fibrosis stages.** Heatmap of top seven DEGs in spots annotated as (A) normal hepatocyte and (B) normal portal tract between early/intermediate and late- stage fibrosis. Venn diagram showing upregulated genes in (C) early/intermediate- and (D) late-stage fibrosis that overlap across DEGs analyses in ballooning, interface, steatosis, and inflammation (portal and lobular) annotation groups. (E) Gene set co-expression scores for the indicated Hallmark gene sets projected onto the UMAP, coloured by Geneset co-expression scores.

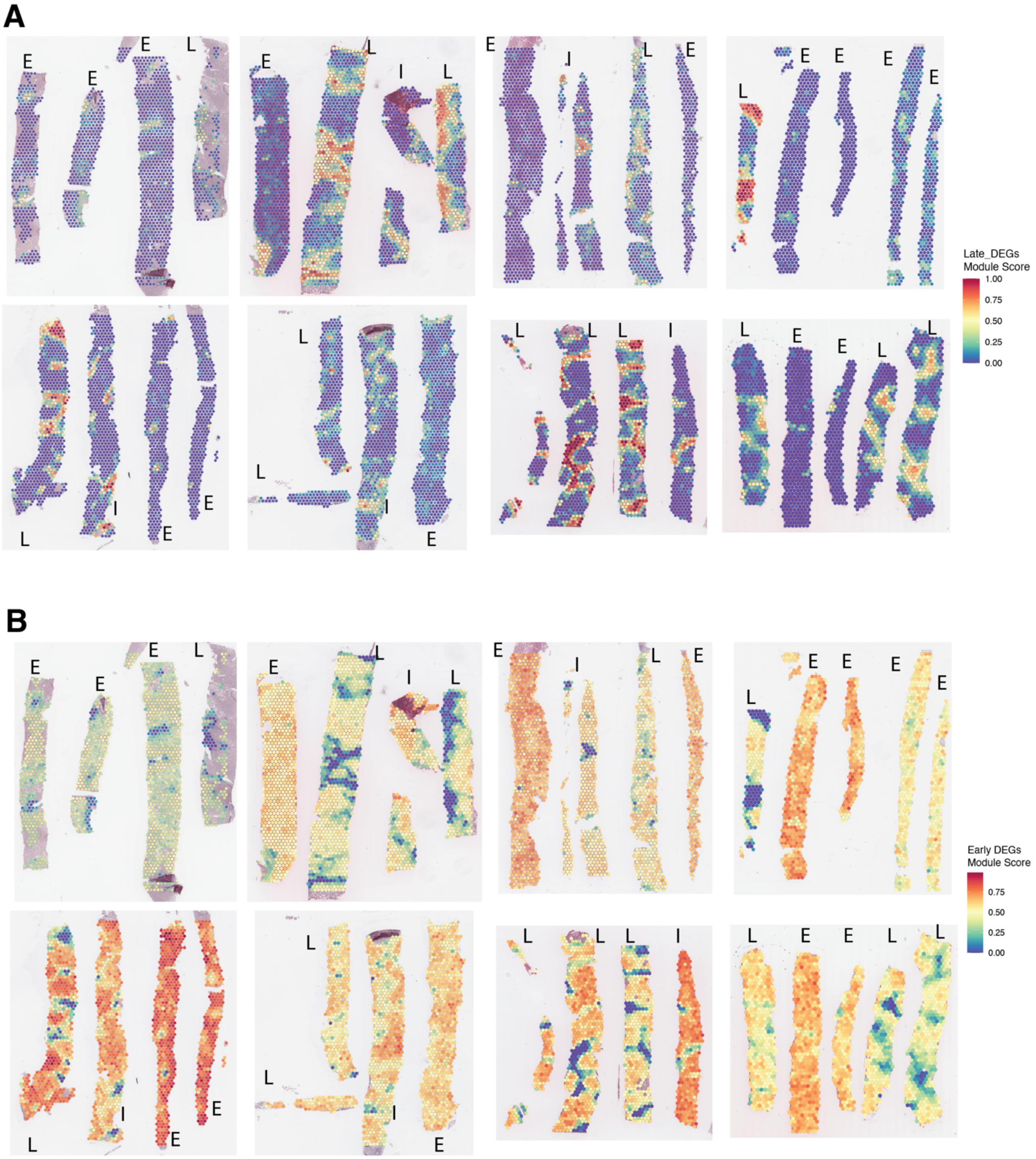


**Figure S4. Spatial distribution of enriched DEGs across liver biopsies.** (A) Late-stage fibrosis enriched differentially expressed genes (DEGs) present in two or more anatomical comparisons between early- and late-stage biopsies, with spots colored by gene expression. (B) Early-stage fibrosis enriched DEGs present in two or more anatomical comparisons, with spots colored by gene expression. Scale bars represent module scores of respective DEG sets.

A

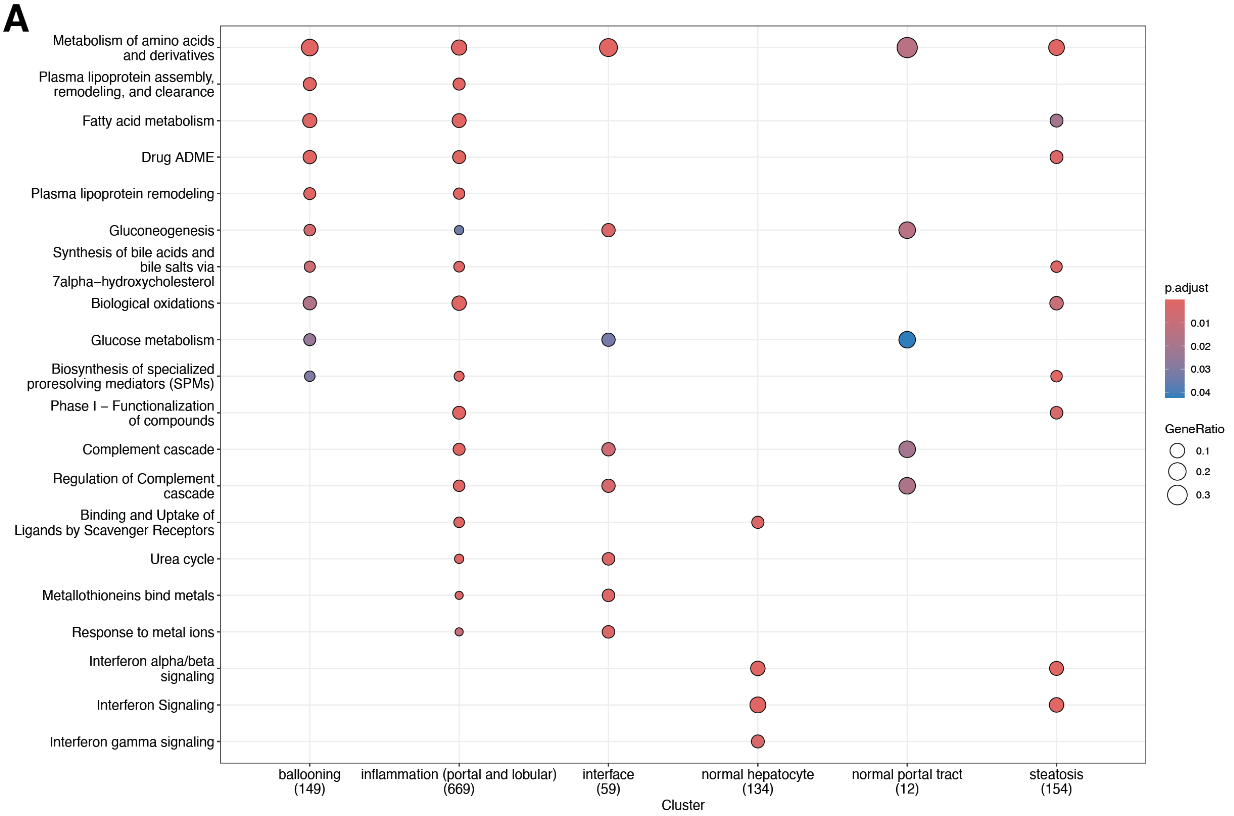
Metabolism of amino acids and derivatives
Plasma lipoprotein assembly, remodeling, and clearance
Fatty acid metabolism
Drug ADME
Plasma lipoprotein remodeling
Gluconeogenesis
Synthesis of bile acids and bile salts via 7alpha−hydroxycholesterol
Biological oxidations
Glucose metabolism
Biosynthesis of specialized proresolving mediators (SPMs)
Phase I − Functionalization of compounds
Complement cascade
Regulation of Complement cascade
Binding and Uptake of Ligands by Scavenger Receptors
Urea cycle
Metallothioneins bind metals
Response to metal ions
Interferon alpha/beta signaling
Interferon Signaling
Interferon gamma signaling
ballooning (149)
inflammation (portal and lobular) (669)
interface (59)
normal hepatocyte (134)
normal portal tract (12)
steatosis (154)
Cluster
p.adjust
0.01
0.02
0.03
0.04
GeneRatio
0.1
0.2
0.3

B

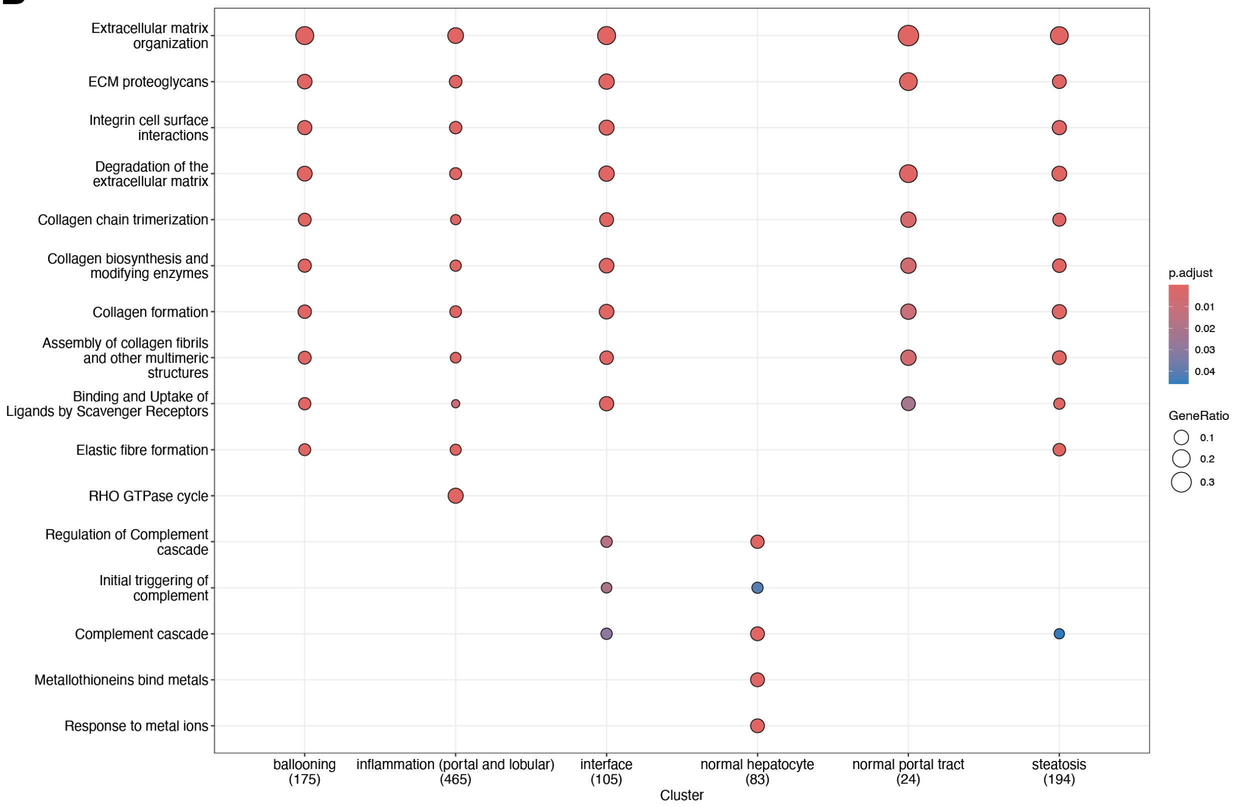
Extracellular matrix organization
ECM proteoglycans
Integrin cell surface interactions
Degradation of the extracellular matrix
Collagen chain trimerization
Collagen biosynthesis and modifying enzymes
Collagen formation
Assembly of collagen fibrils and other multimeric structures
Binding and Uptake of Ligands by Scavenger Receptors
Elastic fibre formation
RHO GTPase cycle
Regulation of Complement cascade
Initial triggering of complement
Complement cascade
Metallothioneins bind metals
Response to metal ions
ballooning (175)
inflammation (portal and lobular) (465)
interface (105)
normal hepatocyte (83)
normal portal tract (24)
steatosis (194)
Cluster
p.adjust
0.01
0.02
0.03
0.04
GeneRatio
0.1
0.2
0.3

**Figure S5. Reactome pathway enrichment analysis of upregulated DEGs across fibrosis stages.** (A) Significantly enriched Reactome pathways for upregulated DEGs in early/intermediate-stage fibrosis, separated per annotation, identified through gene set enrichment analysis. (B) Significantly enriched Reactome pathways for upregulated DEGs in late-stage fibrosis, separated per annotation, identified through gene set enrichment analysis.

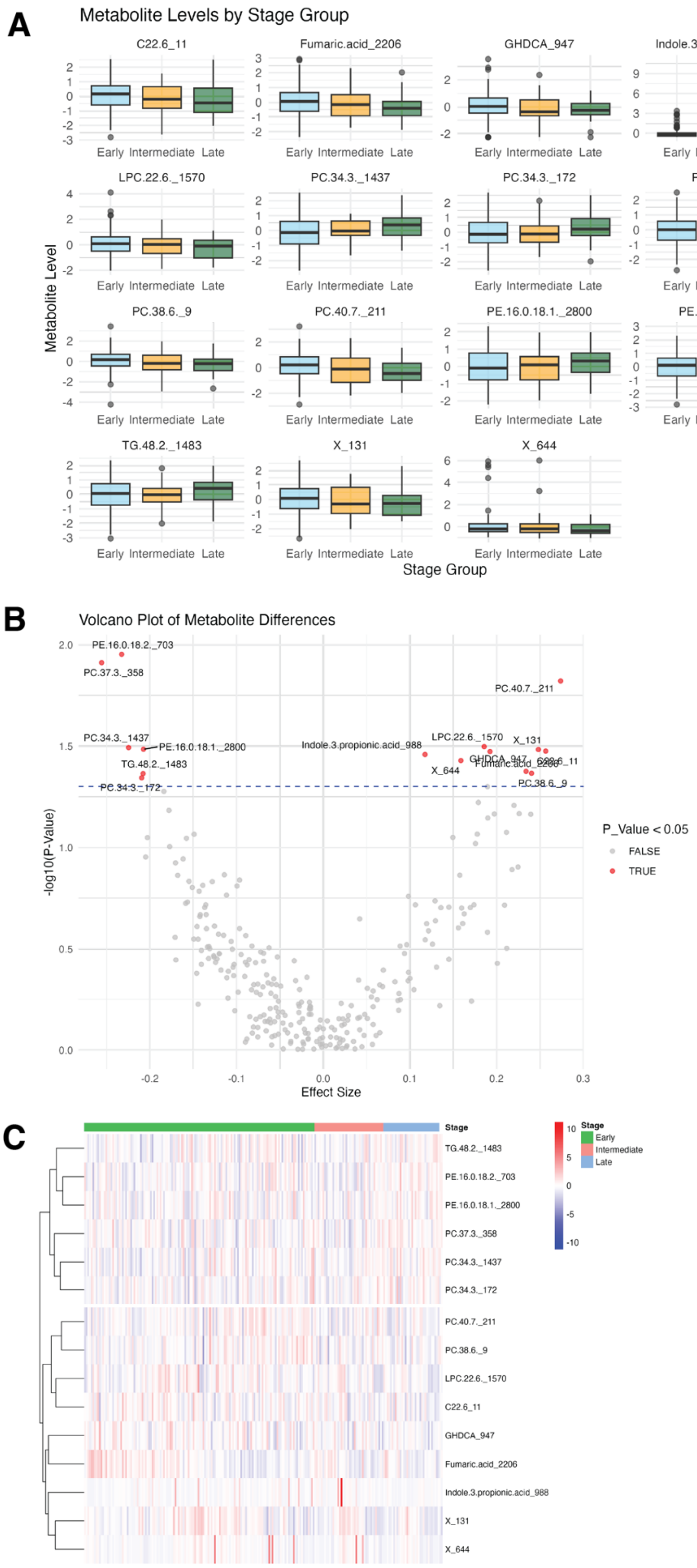

A
Metabolite Levels by Stage Group
C22.6_11
Fumaric.acid_2206
GHDCA_947
Indole.3.propionic.acid_988
LPC.22.6._1570
PC.34.3._1437
PC.34.3._172
PC.37.3._358
PC.38.6._9
PC.40.7._211
PE.16.0.18.1._2800
PE.16.0.18.2._703
TG.48.2._1483
X_131
X_644
Early Intermediate Late
Metabolite Level
Stage Group
B
Volcano Plot of Metabolite Differences
-log10(P-Value)
Effect Size
P_Value < 0.05
FALSE
TRUE
C
Stage
Early
Intermediate
Late

**Figure S6. Differential abundance analysis of serum metabolites between fibrosis stages.** (A) Top 15 metabolites ranked from most to least different between late and early fibrosis stage. (B) Differential abundance of all metabolites with a defined annotation across 216 patients. Red points show metabolites with p-value <0.05 (un-adjusted). (C) Heatmap showing abundance levels of annotated metabolites across all patients (grouped by rows, with the top six rows show metabolites higher in the late stage compared to the early stage, and the remaining rows are lower in the late stage).

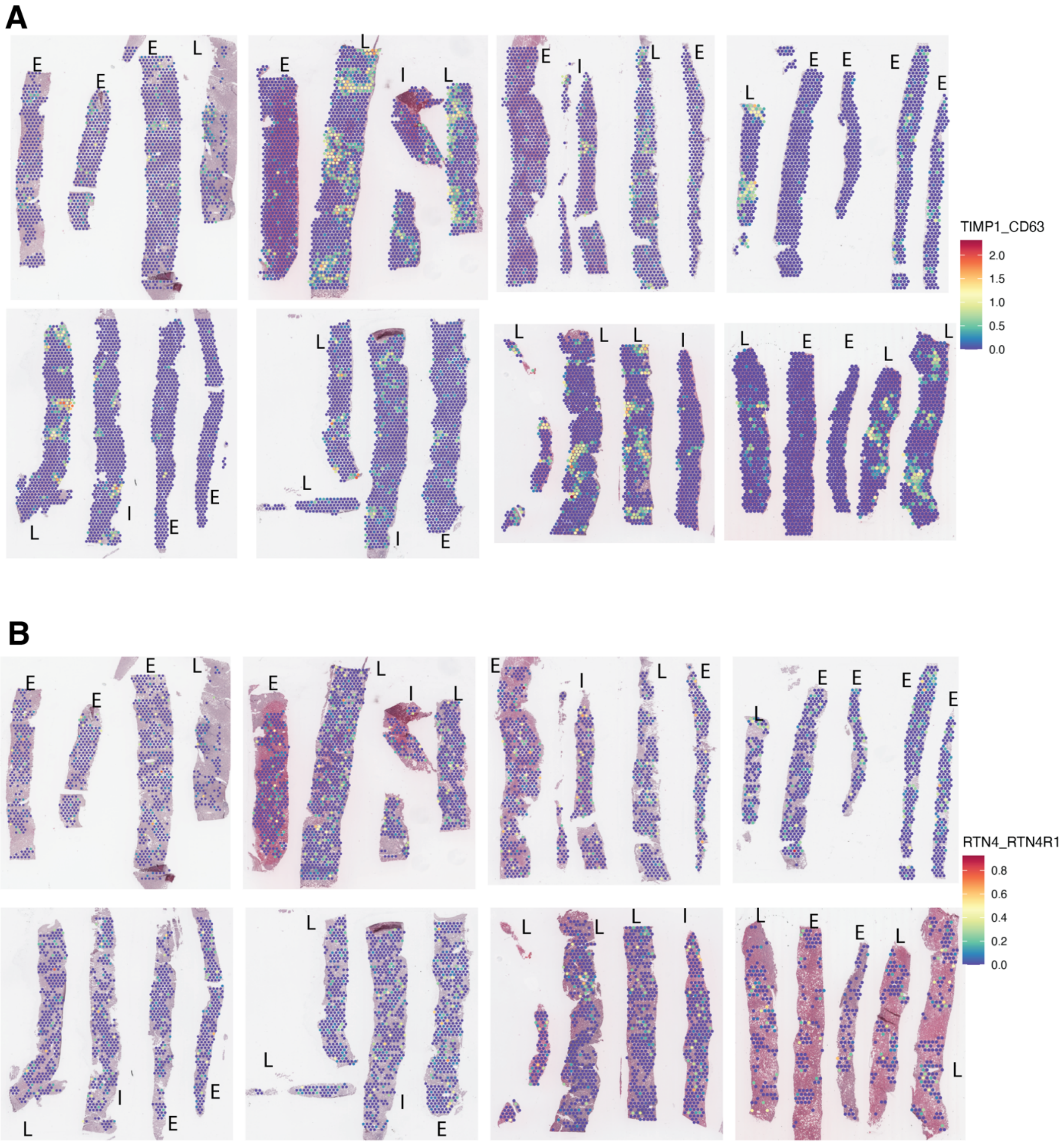


**Figure S7. Spatial visualization of ligand-receptor pairs correlated with fibrosis stage**. Spatial visualisation of the top LR pairs positively (TIMP1-CD63) (A) and negatively (RTN4-RTN4R1) (B) correlated with fibrosis stage across all liver biopsies.
